# A Potent Pandemic Avian Influenza Virus Vaccine Based on a 4^th^ Generation Fully Deleted Adenoviral Vector

**DOI:** 10.1101/2024.12.30.630761

**Authors:** Yan Qi, E. Bart Tarbet, Janae Wheeler Cull, Xianghua Zhang, Uwe D Staerz

**Author notes:** **Correspondence:** Uwe D Staerz, MD, PhD, Research and Development Greffex, Inc., 12635 East Montview Blvd, Aurora, CO 80045, USA E.

## Abstract

The GreVac system was developed as a fast and flexible plug-and-play vaccine platform based on an architecture of fully deleted (fd) helper virus-independent (hi) adenoviral (Ad) vectors. The present study established the potency of the GreVac technology. It demonstrated that the GreFluVie5 vaccine fully protected mice against lethal challenges with the *A/Vietnam/1203/2004* (H5N1) pandemic avian influenza virus. The GreFluVie5 vector delivered a transgene expression cassette for the H5 hemagglutinin and N1 neuraminidase influenza genes. Its fd Ad genome was carried in a capsid of the human serotype 5 (Ad5). The efficacies of three different doses and three different administration routes were compared in the mouse model. The vaccine fully protected animals against viral challenges with the wild-type *A/Vietnam/1203/2004* virus, whose replication in the recipients’ lungs was terminated. It induced strong immune responses. The present experiments also revealed that the *intra muscular* (*i.m.*) delivery route of GreFluVie5 was more efficient than *sub cutaneous* (*s.c.*) or *intra nasal* ones (*i.n.*). Based on results of this animal trail and GreVac’s intrinsic versatility and fast development time, we believe that this platform is ideally suited to swiftly deliver powerful vaccines to infectious diseases with high eruption potentials.

## BACKGROUND

Influenza, commonly also called *the flu*, is an infectious disease of the respiratory tract caused by RNA viruses of the family orthomyxoviridae of the genera A, B and C (Kamps et al., 2006; Krammer et al., 2018). In the case of the influenza A virus a huge reservoir exists in wild birds (Venkatesh et al., 2018). The segmented nature of the influenza genome facilitates genetic re-assortments (*genetic shift*) of existing virus serotypes leading to the emergence of new influenza strains. This may include ones with pandemic potential(Kamps et al., 2006; Krammer et al., 2018; Olsen et al., 2006). Genome mutations (*genetic drift*) drive the appearance of new influenza strains every 1 to 2 years (Parrish et al., 2015). Besides the more benign seasonal outbreaks, virulent pandemic strains appear every 10 to 50 years that may cause millions of deaths (Kamps et al., 2006). More recent highly pathogenic avian influenzas (HPAI) fortunately failed to acquire effective human-to-human transmission. This was exemplified by the lethal *A/Vietnam/1203/2004* H5N1 avian influenza that had erupted in Asia in 2003 through bird-to-human transmissions (He & Kam, 2024; Kamps et al., 2006; Sutton, 2018). It subsequently spread throughout the world with Egypt being its hotspot (Refaey et al., 2015). It never acquired an efficient human-to-human infectivity. However, different studies have now been able to demonstrate that only few mutations boosted its range (Herfst et al., 2012; Imai et al., 2012; Russell et al., 2012). A more widespread pandemic may indeed become reality once the underlying virus acquired enhanced human infectivity. As new influenza variants elude pre-existing human immunity, new vaccines must constantly be developed.

Highly infectious influenzas spread through a human population within time frames that cannot be met by present vaccine development strategies (Buckland, 2015; Lambert & Fauci, 2010). We at Greffex developed the GreVac system as a fast and flexible plug-and-play vaccine platform based on our proprietary architecture of fully deleted (fd) helper virus-independent (hi) adenoviral (Ad) vectors of different serotypes (Lee et al., 2019; Qi et al., 2024). We removed all endogenous Ad genes from the vector genome with the goal to minimize the induction of and interference by anti-Ad responses. We expected that GreVac vaccines better focused the immune system to the vaccine antigen (Abbink et al., 2016; Weaver et al., 2009, 2013) and enabled short-term prime-boost vaccination protocols (Koehler et al., 2006). For its design we did not simply resort to fd Ad vector systems that used helper virus constructs for encapsidation (Cheshenko et al., 2001; D. J. Palmer & NG, 2005; D. Palmer & Ng, 2003). These earlier setups experienced significant contaminations with replication competent adenoviruses (RCA) or helper viruses themselves (Cheshenko et al., 2001; Dormond et al., 2009, 2010). These impurities have the potential to induce potent anti-Ad responses that may in turn interfere with their intended functions (Nomura et al., 2004). GreVac produces fd Ad vectors independently of a helper virus and avoids RCAs (Qi et al., 2024). It was built upon two independently modifiable components, the fully gutted GreVac vector genomes, and the non-packagable circular packaging expression plasmid. We found that GreVac was able to provide the basis for numerous vaccines (*data not shown*) and to deliver a vector with a new transgene expression cassette within one month (Qi et al., 2024). As the GreVac scheme incorporates versatility and speed, it may ultimately change the present paradigm of immune prophylaxis from the stockpiling of vaccines against infections with high eruption potentials to a just-in-time production of specific vaccines.

## RESULTS

### Composition of the GreFluVie5 Vaccine

The GreVac-based genome of GreFluVie5 was assembled in a pBR322-derived plasmid (**Figure 1**) (Qi et al., 2024). It could be released from the cloning vector by a simple restriction enzyme cut. It carried a bicistronic expression cassette for the hemagglutinin 5 (H5) (AY818135.1) and neuraminidase 1 (N1) (AY651447.1) of the *A/Vietnam/1203/2004* influenza virus. It was flanked by inverted terminal repeats (ITR) and a packaging signal Y both derived from an Ad of the human serotype 5. To increase the genome to a packageable size, a DNA stuffer was added that consisted of an internal fragment of the human 5-aminoimidazole-4-carboxamide ribonucleotide formyl-transferase/IMP cyclohydrolase (ATIC) gene. Once released from the cloning plasmid, the linear GreFluVie genome was packaged into an Ad capsid of the human serotype 5. The linear genome and a packaging expression plasmid were co-transfected into a variant of the human embryonic kidney (HEK293) cells (Qi et al., 2024). The completed GreFluVie5 vaccine vector was purified by a two-step column chromatography into a vector formulation buffer (VFB). Its functionality was examined by transducing test cells and examining their expression of the transgenes (Eglon et al., 2009).

**Figure 1.**
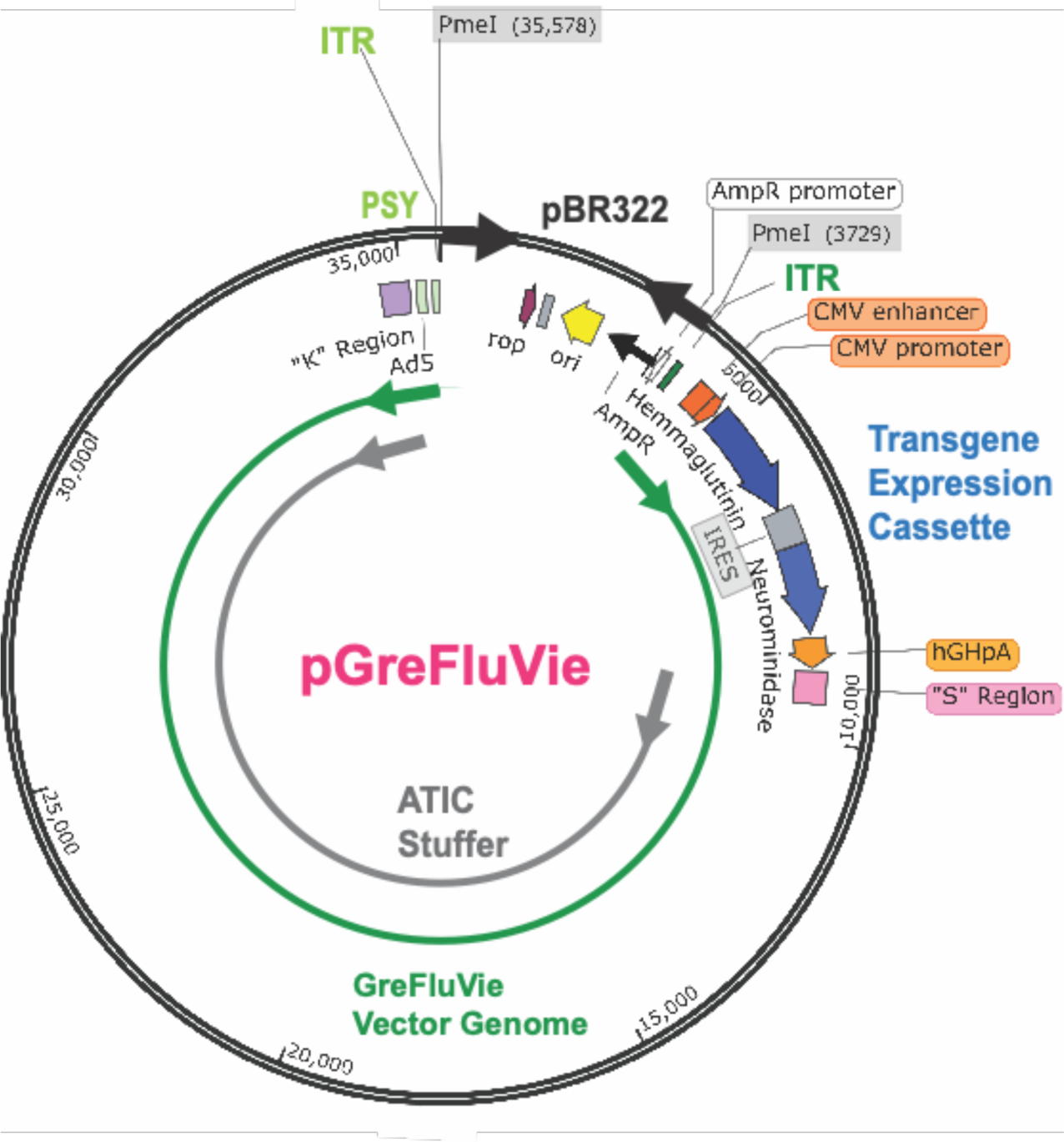
The GreFluVie5 Genome. pGreFluVie carries the genome of the corresponding GreVac vector in a pBR322-based plasmid. Ad5: Adenovirus human serotype 5; AmpR: ampillin resistance gene; ATIC: human 5-aminoimidazole-4-carboxamide ribonucleotide formyl-transferase/IMP cyclohydrolase; CMV: cytomegaly virus; hGH: human growth hormone; S: S cloning region; ITR: inverted terminal repeat; K: K cloning region: ori: origin of replication; PmeI: restriction enzyme; PSY: Adenovirus packaging site; rop: repressor of primer.

### Mouse Challenge Study

The objective of the present study was to establish the efficacy of the GreFluVie5 vaccine to protect mice against a lethal challenge with the pandemic avian influenza strain *A/Vietnam/1203/2004* (H5N1). The mice were distributed into 11 experimental groups (**Table 1**). The optimal doses and routes of administration were determined, and prime-boost vaccination protocols were examined (*illustrated in* **Figure 2**). The efficacies of *i.m*., *s.c. and i.n.* delivery routes were compared with three different vaccine doses, which consisted of 3 x 10^8^ genome equivalents (GE) (**High Dose**), 3 x 10^7^ (GE) (**Medium Dose**) and 3 x 10^6^ GE (**Low Dose**) per immunization. The **Positive Control** vaccine contained approximately 5 μg total hemagglutinin plus 0.2% Alum and was administered *i.m.* The **Negative Control** consisted of the VFB. We studied both immune responses and protections against the influenza virus infection. Three weeks after the last vaccine dose had been administered, the mice received the wild-type *A/Vietnam/1203/2004* virus given via an *i.n.* instillation. To examine immune responses, the animals were bled twice, prior to the vaccine boost and prior to the virus infection. Post-challenge, they underwent lung lavages on days 4 and 7 thereafter to determine lung virus titers and secreted IgA (sIgA) levels, and they were weighed initially daily and then every second day.

**Figure 2.**
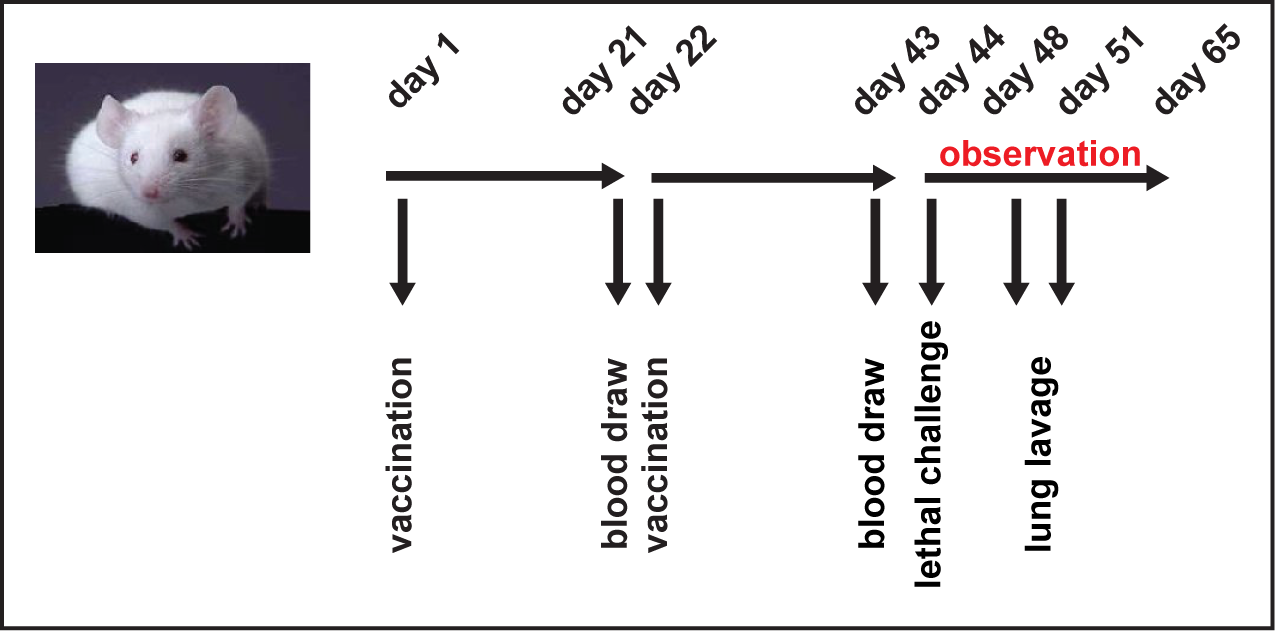
Mouse Lethal Challenge Study. Balb/c mice were bled on days 21 and day 43 and vaccinated on days 1 and 22. They received the challenge virus *A/Vietnam/1203/2004* on day 44. They were observed until day 65.

### Health Status

Following the viral challenge, the animals were observed for weight loss and mortality through day 21. **Figure 3** shows Kaplan-Meier survival curves and mean body weight changes for the different groups of vaccinated and control mice. The **Negative Controls** suffered greatly. They showed significant body weight losses, and almost all of them succumbed to the virus infection by day 17. All **High Dose** mice were completely protected when they had received the vaccine via the *i.m.* or *s.c.* routes. They only showed minor losses in body weight (< 3%) when compared to other mice at the same facility that were not involved in the trial. At this dose, the GreFluVie5 efficacy exceeded the 91% survival seen for the **Positive Controls**, whose average body losses reached approximately 12%. Vaccinations via the *i.n.* route proved less efficient. At the **High Dose** 18% of mice succumbed to the influenza. Moving to the **Medium Dose** reduced the overall protection provided by GreFluVie5. Yet, 91% of the animals survived, when this vaccine dose was given as an *i.m.* injection. These mice still fared better than the **Positive Control** ones. On average they only experienced less than 8% in body weight loss. At this **Medium Dose**, protection fell to 58% when the GreFluVie5 was given via *s.c.* injection and faded to 16% for the *i.n.* delivery route. In both cases, the animals were not able to maintain their body weight. At the **Small Dose**, this vaccine proved inefficient at all immunization route.

**Figure 3.**
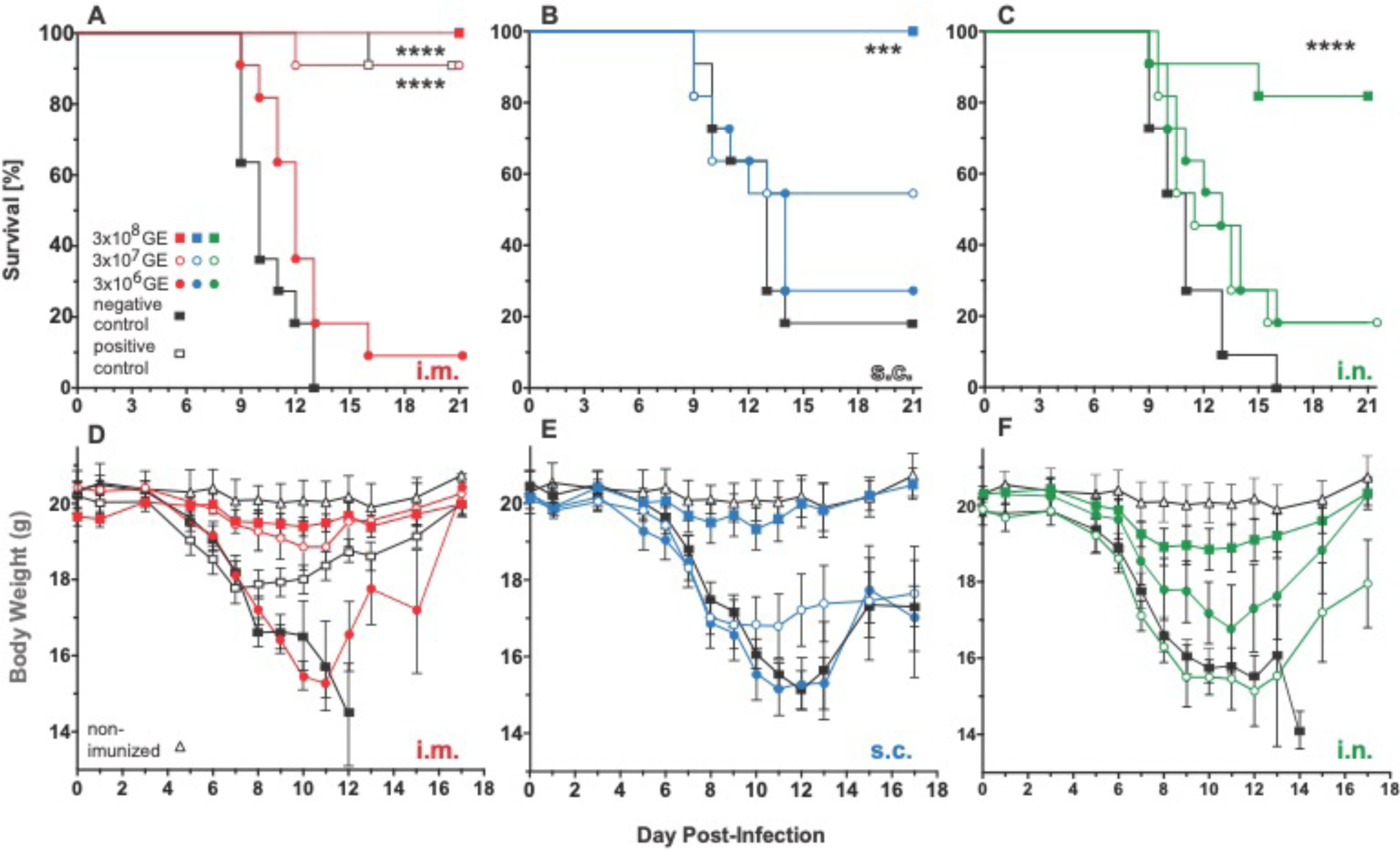
Survival and Body Weight After Viral Challenge. Mice received the 3 x 10^8^ GE (**High Dose**: 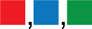), 3 x 10^7^ GE (**Medium Dose**: 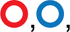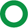), 3 x 10^6^ (**Low Dose**: 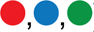) of the GreFluVie5 vaccine, the hemagglutinin protein vaccine (**Positive ∗ Control: ∗∗ □**) or the ∗∗∗ Vector ∗∗∗∗ Formulation Buffer (▀) by an *i.m.* (**A, D**), *i.n.* (**B, E**) or *s.c.* (**C, F**) delivery. They were surveilled for health and their body weights were determined. : p ≤ 0.05; : p ≤ 0.01; : p ≤ 0.001; : p ≤ 0.0001.

### Virus Lung Titers

On day 4 and day 7 post-challenge, some mice of each group underwent lung lavages. The retrieved samples were tested for the presence of the live *A/Vietnam/1203/2004* virus. As depicted in **Figure 4, Negative Controls** showed high virus loads at both time points. The **Positive Control** vaccine failed to significantly lessen the virus burden. When **High Doses** of GreFluVie5 were administered as *i.m.* or *s.c.* injections, the lung virus titers were substantially decreased as early as day 4. They became undetectable in most animals of these groups by day 7. Decreasing the vaccine dose to **Medium** or **Low** abolished anu virus titer reduction. Although the *i.n.* delivery of the vaccine resulted in undetectable virus titers in in 2 of 5 of the **High Dose** animals, this reduction failed to reach statistical significance.

**Figure 4.**
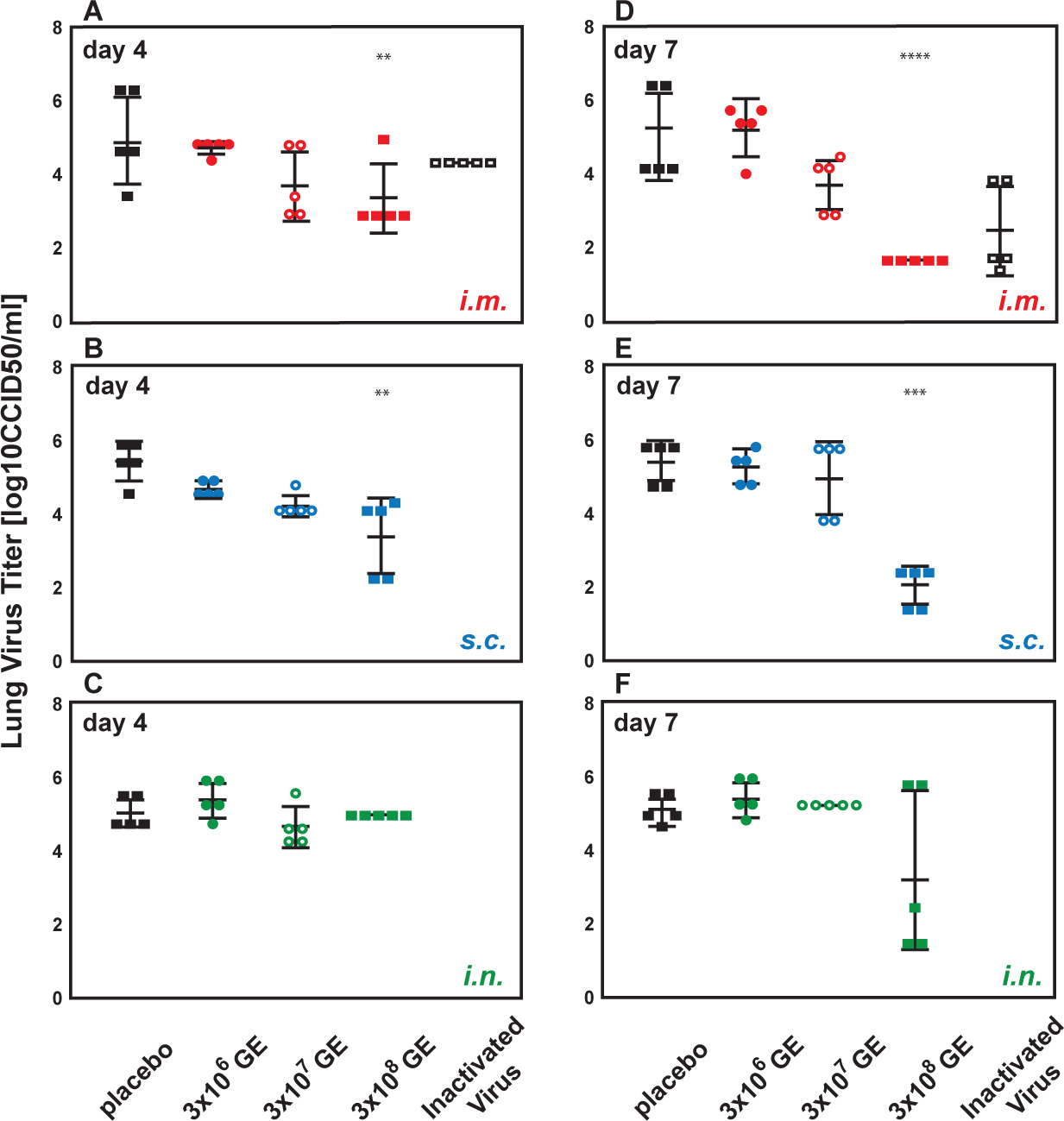
Lung Virus Titers. On days 4 (**A, B, C**) and 7 (**D, E, F**) post challenge, the mice that had been immunized *i.m.* (**A, D**), *s.c.* (**B, E**) or *i.n.* (**C, F**) with GreFluVie6, or had received the **Positive Control** vaccine (**A, D**) or the **VFB** (**A, B, C, D, E, F**) underwent lung lavages. The retrieved material was examined for the presence of live *A/Vietnam/1203/2004* viruses (*see* Figure 3 for symbols and p values)

### Immune Responses

All animals were bled after the primary and secondary immunizations (**Figure 2**). Levels of *A/Vietnam/1203/2004*-neutralizing antibodies were determined. As seen in **Figure 5**, mice of all vaccine groups raised significant antibody responses already after the first vaccine dose. This included all mice that had received the GreFluVie5 vaccine at any tested dose or administration route. Antibody titers increased with the vaccine dose. In addition, the second immunization boosted the antibody titers in all cases. No significant differences in antibody responses were observed for the different vaccination routes. The seroconversion rate was 100% for all doses and all delivery venues.

**Figure 5.**
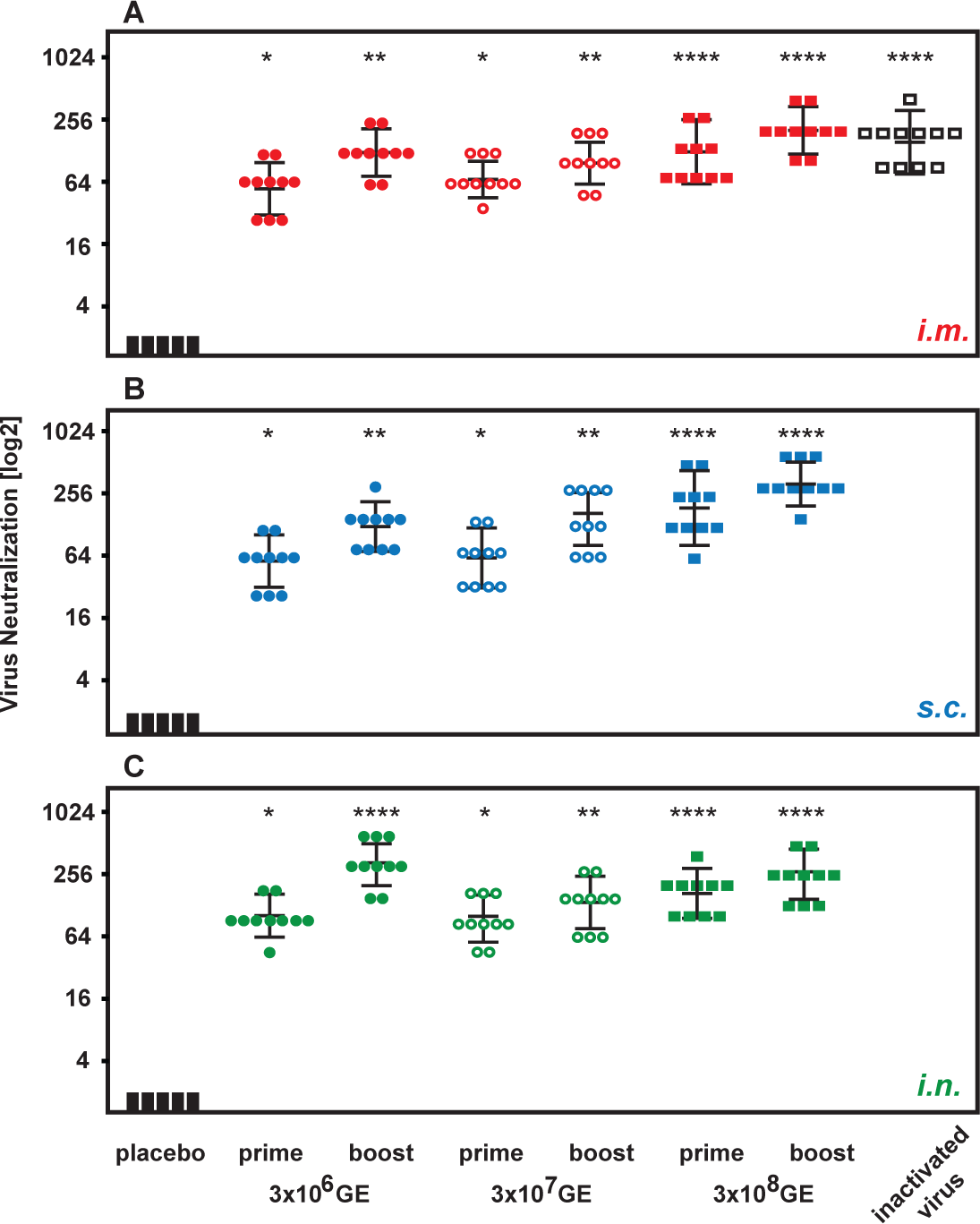
Neutralizing Antibodies. Mice that been vaccinated once (day 1) or twice (day 22) with the GreFluVie6 vaccine via the *i.m.* (**A**), *s.c.* (**B**) or *i.n.* (**C**) route, or had received the **Positive Control** vaccine (**A**) or the VFB (**A, B, C**) were bled. Virus neutralizing antibody titers were determined in their sera (*see* Figure 3 for symbols and p values).

A similar picture was seen when the specific antibody levels were determined in a hemagglutination inhibition assay (**Figure 6**). In this assay, the immune responses were more sensitive to the vaccine dose. Whereas **Low Dose** mice showed lesser antibody titers for all delivery routes, the results for **High Dose** animals remained at the same level. Second booster effects seemed less pronounced.

**Figure 6.**
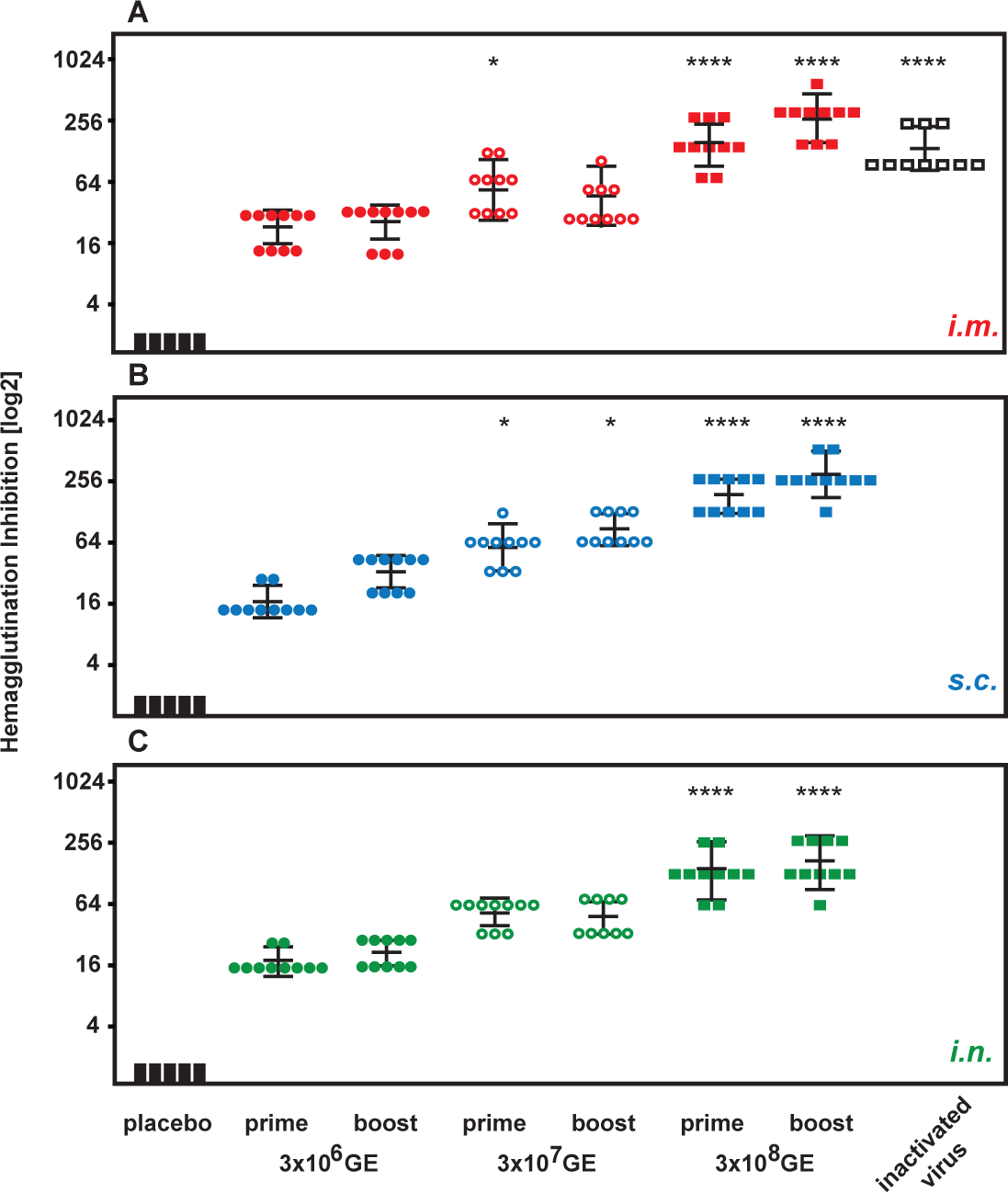
Neutralizing Antibodies. Mice that been vaccinated once (day 1) or twice (day 22) with the GreFluVie6 vaccine via the *i.m.* (**A**), *s.c.* (**B**) or *i.n.* (**C**) route, or had received the **Positive Control** vaccine (**A**) or the VFB (**A, B, C**) were bled. Virus red blood cell hemagglutination inhibiting antibody titers were determined in their sera (*see* Figure 3 for symbols and p values).

Evaluations of the immune responses following the vaccinations also included the measurements of sIgA levels in lung lavage samples. On day 4 post-challenge, **Negative Control** mice presented with low sIgA levels (**Figure 7**). Elevated sIgA concentrations were detected in all **High Dose** animals. However, this rise was more pronounced when the vaccine was administered via *s.c.* and *i.n.* routes rather than an *i.m.* injection. At the **Low Dose**, only the *i.n.* administration showed significant increased sIgA levels in the lung. On day 7, the sIgA concentrations surged in the **Negative Control** animals. This upturn was most probably due to the intrinsic immune responses that were induced by the viral infection. At this point, significantly lowered sIgA levels were not observed for any of the immunized animals.

**Figure 7.**
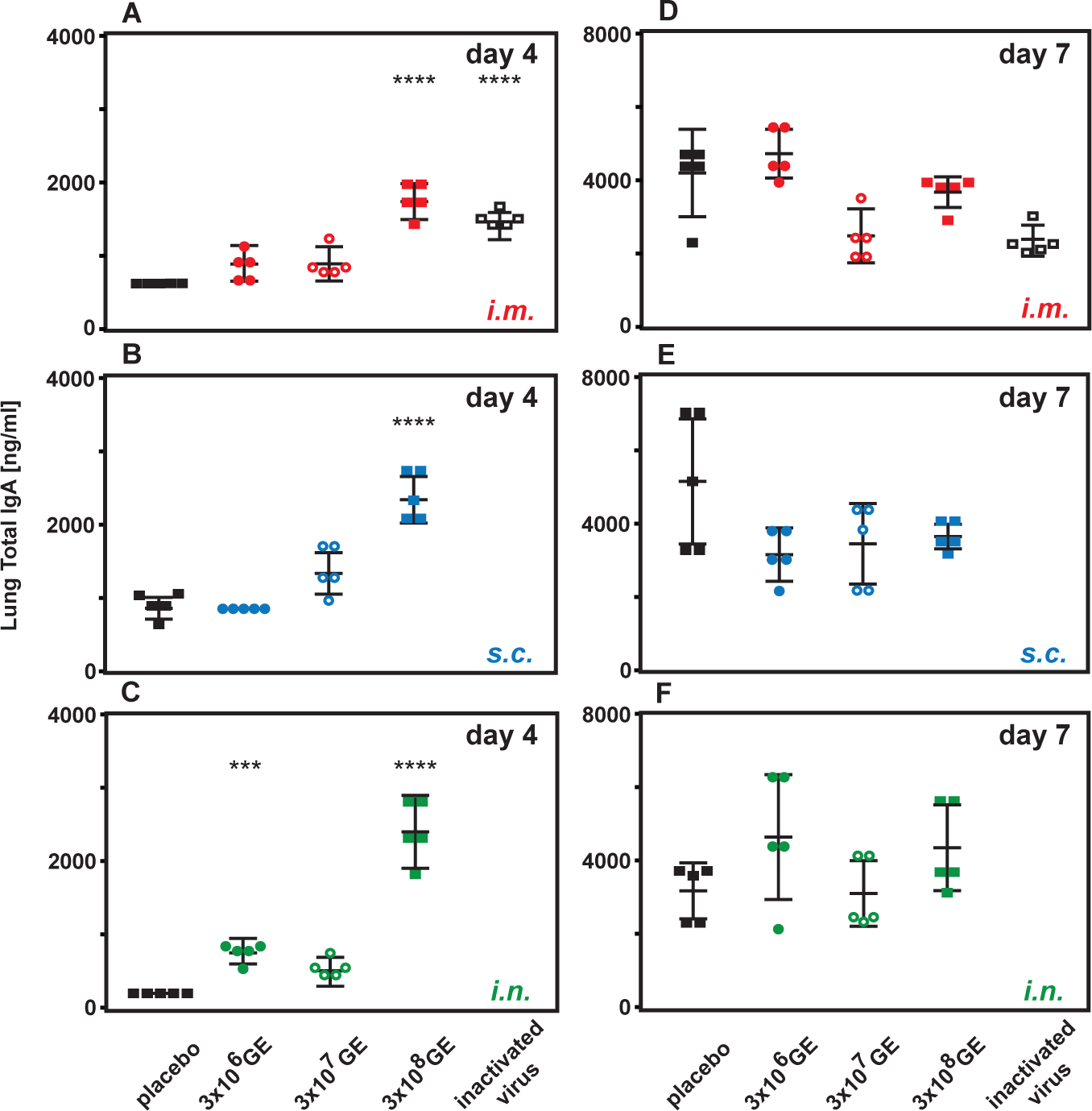
sIgA Levels in the Lung. On days 4 (**A, B, C**) and 7 (**D, E, F**) post challenge, the mice that had been immunized *i.m.* (**A, D**), *s.c.* (**B, E**) or *i.n.* (**C, F**) with GreFluVie6, or had received the **Positive Control** vaccine (**A, D**) or the VFB (**A, B, C, D, E, F**) underwent lung lavages. The retrieved material was examined for the presence of secreted IgA (sIgA) levels (*see* Figure 3 for symbols and p values).

## DISCUSSION

The present study aimed to demonstrate that the GreVac approach represented a technology platform to develop potent vaccines. GreVac uses an advanced strategy to produce fully deleted Ad vectors without the participation of a helper virus (Qi et al., 2024). In its GreFluVie5 iteration, the Ad genome carried a transgene expression cassette for the H5 hemagglutinin and the N1 neuraminidase (**Figure 1**). The potency of this vaccine was evaluated in a mouse model, in which protection against infections with an non-animal-adapted wild-type *A/Vietnam/1203/2004* pandemic influenza virus could be examined (**Figure 2**, **Table 1**).

The efficacies of different vaccine doses and administration routes were compared. After prime/boost immunizations, the animals were exposed to the wilde-type *A/Vietnam/1203/2004* virus. The viral challenges were provided through *i.n.* instillations to mirror the natural infection paths of an influenza virus. Their health status, here body weights, and survivals were observed through day 21 (**Figure 3**). Prior to this study, we had assumed that an *i.n.* vaccination provided the best defense against a mucosal infection, such as a viral influenza. Although the GreFluVie5 vaccine raised strong responses independently of the delivery route, the *i.m.* injection proved most efficient. It delivered full protection when given at the **High Dose** level (**Figure 3**). It still shielded more than 90% of animals to the level of the **Positive Control**, even after its dose was reduced by ten-fold to **Medium**. At this level protection became more limited when given via the *s.c.* route and faded when an *i.n.* delivery was used. Effectively vaccinated animals not only survived, but also remained in good health. Animals that had received the GreFluVie5 vaccine at the **High Dose** faired best. Only minor losses in body weight were observed when compared to normal mice maintained in parallel at the same facility.

Viral pneumonias are one of the most dangerous complications of influenza infections (Gomez Lorenzo & Fenton, 2013). We wanted to know whether GreFluVie5 affected the presence of the *A/Vietnam/1203/2005* influenza virus in the lung. The present results demonstrated that it was possible to fully remove active virus from the lung tissue (**Figure 4**). This was achieved when the vaccine was given at the **High Dose** level as an *i.m.* injection. Our original idea that a vaccine delivery to the respiratory tract, *i.e.* an *i.n.* immunization, would be best suited to protect the lung, proved incorrect. As our vaccine suppressed the virus levels to undetectable by day 7, it could be concluded that it reached the ultimate goal of vaccination, *i.e.* infectious sterility.

The vaccine’s protective function was mirrored in its immunogenicity. Animals were bled. Virus neutralization assays with the wild-type *A/Vietnam/1203/2004* were used to as one measure of humoral immune responses (**Figure 5**). The first vaccination with the **Low Dose** already induced strong humoral responses in all animals whether administered via *i.m.*, *s.c.* or *i.n.* routes. Increasing the vaccine dose stepwise further increased the antibody titers. A second vaccination further boosted the immune responses in all animals. Hemagglutination inhibition assays as a second read-out of humoral immunity drew a similar picture (**Figure 6**). Broad immune responsiveness to all vaccine doses and delivery routes. However, using this read-out, the dose-response curves were steeper.

As surrogate measures of immune responses in the lung, we measured sIgA levels in the lung lavage samples. On day 4 post-challenge, the material retrieved **Negative Control** mice only had low sIgA concentrations (**Figure 7**). We concluded that at this point the animals’ immune systems had not responded to the infectious insult. High total sIgA concentrations in the lung lavage material measured after vaccination would therefore represent evidence of a successful immunization. Indeed, all mice that had received the vaccine at the **High Dose** presented with elevated sIgA levels. Even though the sIgA concentrations seemed highest after an *i.n.* delivery, the differences with the other administration routes were not significant. On day 7 post-challenge, the sIgA levels of the **Negative Control** as well as the one of the other animals reached high levels. At this point, we concluded that immune responses raised by the viral infection superseded the vaccine induced ones.

GreVac-based vectors are completely gutted of their Ad genes. They represent a 4^th^ generation of Ad vectors that are produced in the absence of and without the contaminations with helper viruses (Dormond et al., 2009, 2010). The only Ad-derived antigens are found as proteins of the Ad particle. In earlier studies, we had observed that GreVac-based vectors only induced weak anti-Ad immunity (Qi et al., 2024). We had therefore hypothesized that such responses would not interfere with the effectiveness of a second dose. The present results indeed proved that GreFluVie5, a GreVac vaccine, could be successfully used in short-order prime/boost vaccination protocols. A change of vaccine design is no longer necessary for a second dose.

In summary, these studies demonstrated that a GreVac-based vaccine induced strong immune responses and provided potent protection against a lethal challenge with the wild-type *A/Vietnam/1203/2004* virus. They also revealed that the *i.m.* delivery route proved more efficient than *s.c.* or *i.n.* ones not only for the induction of immune responses, but also for the suppression of virus replication in the lung. This finding was unexpected as others had described the superiority of mucosal vaccinations (Casimiro et al., 2003; Chen et al., 2024; Coughlan et al., 2018; Happe et al., 2024; Jia et al., 2024; Jin et al., 2022; Li et al., 2023; Tapia et al., 2016)Based on the observed dose-responses, we calculated that the *i.m.* vaccination was about 30-fold more efficient than an *i.n.* one. Furthermore, deleting all endogenous Ad genes from the GreFluVie5 genome allowed effective short-order prime/boost immunization protocols. In several earlier reports, early generation Ad vectored vaccine had used a completely different design for a second dose (Casimiro et al., 2003; Chen et al., 2024; Coughlan et al., 2018; Happe et al., 2024; Jia et al., 2024; Jin et al., 2022; Li et al., 2023; Tapia et al., 2016) The GreFluVie5 doses that were used in this study were significantly lower than the ones that had been reported with vaccine based on early generation Ad vectors (Gao et al., 2006; Morse et al., 2013; Patel et al., 2010; Singh et al., 2008). This finding supported our idea that fully deleted Ad vaccine better targeted the immune system and more efficiently induced immune protection.

## MATERIAL AND METHODS

### Animals

Female 6 week-old BALB/c mice were obtained from *Charles River Laboratories*. The mice were quarantined for 72 hours before use. They were maintained on Teklad Rodent Diet (*Harlan Teklad*) and tap water at the Laboratory Animal Research Center of Utah State University (USU).

### Challenge Virus

The influenza virus A*/Vietnam/1203/2004* (H5N1) was obtained from the Centers for Disease Control (Atlanta, GA). Viral propagation was done in Madin-Darby canine kidney (MDCK) cells (*American Type Culture Collection*). Parent virus was passaged once to prepare a challenge pool. The challenge pool was then titrated in MDCK cells before use. The cells were grown in MEM containing 5% fetal bovine serum (*Hyclone, Logan, UT*) with no antibiotics in a 5% CO2 incubator.

### Vaccine

GreFluVie6 (H5N1) vaccine was produced and purified as described in (Qi et al., 2024). It was diluted at concentrations of 7.5 x 10^9^ GE/ml (**High Dose**), 7.5 x 10^8^ GE/ml (**Medium Dose**) or 7.5 x 10^7^ GE/ml (**Low Dose**) suspended in vector formulation buffer (VFB, Tris HCl 20mM pH7.8, NaCl 140mM, MgCl_2_ 5mM, EDTA 0.5mM, polyethylene glycol 4000 (PEG 4000) 0.02 mM, and sucrose 2% (W/V)). The **Positive Control** vaccine was composed of a killed whole virus vaccine produced at USU that consisted of the influenza *A/Vietnam/1203/2004 x Ann Arbor/6/60* hybrid virus grown in MDCK cells. After the virus was harvested from MDCK cells, it was inactivated by addition of binary ethyleneimine (BEI) and then clarified/concentrated using tangential flow filtration using a Pellicon® XL 50 cassette (100kd, *Millipore*). The dose of USU vaccine that would protect 100% of mice from influenza *A/Vietnam/1203/2004* (H5N1) virus challenge dose had been determined in a previous study.

### Animal Studies

Animal numbers and study groups are described in **Table 1**. Groups of mice (12 per group) were vaccinated by the *i.m.*, *s.c.* or *i.n.* routes in 40 μl of volume on two occasions, day 1 and 22, with each vaccine dose containing 3 x 10^8^ GE (**High Dose**), 3 x 10^7^ GE (**Medium Dose**), or 3 x 10^6^ GE (**Low Dose**) of the GreFluVie5 vaccine. The **Placebo Groups** received 40 μl of VFB by the same routes on the same vaccination schedules. The **Positive Control** vaccine contained approximately 5 μg total hemagglutinin (*USU*) plus 0.2% Alum (Alhydrogel®) and was administered in a 50 μl volume on a single occasion by the *i.m.* route on day 21 (same day as the second vaccination with the GreFluVie5 vaccine). For influenza virus challenges, groups of mice were anesthetized by intraperitoneal injection of ketamine/xylazine (50 mg/kg//5 mg/kg) prior to *i.n.* challenge with a 90-μl suspension of influenza *A/Vietnam/1203/2004* corresponding approximately 5 plaque forming units (1x LD90) of virus per mouse. All mice were administered virus challenge on study day 44. Mice were weighed prior to virus challenge and then daily or every other day thereafter to assess the effects of vaccination on ameliorating weight loss due to virus infection. All mice were observed for morbidity and mortality through day 21 post-challenge.

### Anti-Influenza Virus Neutralizing Antibody Assay

MDCK cells were seeded in 96-well plates at 1 x 10^4^ cells per well in MEM containing 5% FBS (*Hyclone*) 24 hours prior to use. On the next day, serial 2-fold dilutions of serum samples from five mice were prepared in serum-free media, containing 10 units/ml trypsin and 1 μg/ml EDTA, starting at 1:32 dilution and ending at 1:4096. Each serum dilution was mixed 1:1 (0.1 ml) with serum-free media (containing trypsin and EDTA) containing approximately 100 CCID50/well of the influenza *A/Vietnam/1203/2004 x Ann Arbor/6/60* hybrid virus (Vietnam H5 and N1 surface proteins and Ann Arbor core). After incubation at room temperature for 1 h, the serum-influenza virus mixture (0.2 ml) was transferred to a well containing MDCK cells and incubated for 3 days. Anti-influenza virus neutralizing (VN) antibodies were measured as cytopathic effect (CPE) inhibition. CPE was scored from duplicate samples by examining the MDCK cell monolayers under a light microscope on day 3 post-infection.

### Hemagglutination Inhibition (HAI) Test

Prior of the HAI assay, sera were pre-treated with receptor-destroying enzyme II (RDE; Vibrio cholerae neuraminidase; *Accurate Chemical and Scientific*) to remove non-specific inhibitors by diluting one part serum with three parts enzyme and incubating at 37°C for 18h. RDE was subsequently inactivated by heating in a 56°C water bath for 45 min. Serum samples were diluted in PBS in 96-well round-bottom microtiter plates (*Fisher Scientific*). Following dilution of serum, 8 HA units/well of influenza *A/Vietnam/1203/2004 x Ann Arbor/6/60* hybrid virus (Vietnam H5 and N1 surface proteins and Ann Arbor core) plus turkey red blood cells (*Lampire Biological Laboratories*) were added (50 μl of each per well), mixed briefly, and incubated for 60 min at room temperature. The HAI titers of serum samples are reported as the reciprocal of the highest serum dilution at which hemagglutination was completely inhibited.

### Bronchioalveolar Lavage (BAL)

Animals were killed, lavage procedure was completed within 5 to 10 min of each animal’s death. A volume of 0.75 ml of phosphate buffered saline (PBS) was slowly delivered into the lung through the tracheal tube. Immediately after delivery the fluid was slowly withdrawn by gentle suction and the samples stored at -80°C. The procedure was repeated a total of three times and lavage fluids from each mouse were pooled.

### Lung Virus Titer Determination

BAL samples were centrifuged at 2000 x g for 5 minutes. Varying 10-fold dilutions of BAL supernatants were assayed in triplicate for infectious virus in MDCK cells, with virus titers calculated as described previously (Tao et al., 2017). Virus titers less than 200 were considered negative.

Total IgA levels in lung lavage samples from mice were determined by use of the mouse IgA enzyme immunoassay (EIA) kit (*Bethyl Laboratories, Montgomery, TX*) according to the manufacturer’s instructions. Briefly, goat anti-mouse IgA bound to microtiter plates (*Fisher Scientific*) was used to capture antibody from lavage fluid samples for 1 h at room temperature, after which goat anti-mouse IgA conjugated to horseradish peroxidase was used to detect bound antibody. Antibody concentrations were read off a standard curve generated by using pooled mouse sera calibrated for IgA antibody (*Bethyl Laboratories*).

### Statistical Analysis

Kaplan-Meier survival curves were generated and compared by the Log-rank (Mantel-Cox) test followed by pairwise comparison using the Gehan-Breslow-Wilcoxon test in Prism 6.0f (*GraphPad Software Inc., La Jolla, CA*). The mean body weights were analyzed by analysis of variance (ANOVA) followed by Tukey’s multiple comparison test using Prism 6.0f. Virus titer differences were evaluated by ANOVA on log-transformed values assuming equal variance and normal distribution. Following ANOVA, individual treatment values were compared to placebo control by Tukey’s pair-wise comparison test using Prism 6.0f. The results from virus neutralization assays and hemagglutination inhibition assays were analyzed by ANOVA followed by Tukey’s multiple comparison test using Prism 6.0f.

### Ethics Regulation of Laboratory Animals

This study was conducted in accordance with the approval of the Institutional Animal Care and Use Committee of Utah State University. The work was done in the AAALAC-accredited Laboratory Animal Research Center of Utah State University. The U. S. Government (National Institutes of Health) approval was maintained (Animal Welfare Assurance no. A3801-01) in accordance to the latest National Institutes of Health Guide for the Care and Use of Laboratory Animals.

## Notes

### Competing Interest Statement

The authors, Qi, Cull, Zhang and Staerz are employees of Greffex, Inc. In addition to their salaries they have received stock options as compensation.

